# Osmoregulation affects elimination of microplastics in fish in freshwater and marine environments

**DOI:** 10.1101/2024.10.09.617352

**Authors:** Hilda Mardiana Pratiwi, Toshiyuki Takagi, Suhaila Rusni, Koji Inoue

**Author notes:** To whom correspondence should be addressed. Corresponding Hilda Mardiana Pratiwi, *E-mail address. Abbreviations: Dph, Days post-hatching; FW, Freshwater; GIT, Gastrointestinal tract; GRT, Gut Retention Time; MP, Microplastic; PS, Polystyrene; SW, Seawater.

## Abstract

In recent decades, microplastics (MPs) have emerged as one of the biggest environmental challenges in aquatic environments. Ingestion and toxicity of MPs in seawater (SW) and freshwater (FW) fish have been studied extensively both in field and laboratory settings. However, the basic mechanism of how fish deal with MPs in SW and FW remains unclear, although physiological conditions of fish differ significantly in the two environments. In this study, using Javanese medaka (*Oryzias javanicus*), a euryhaline fish which adapts readily to both SW and FW, we investigated elimination of MPs in fish in SW and FW environments. We exposed *O. javanicus* larvae (21 days post-hatching) to 0.25 mg/L of fluorescent polystyrene microspheres (1 µm) for 24 h and then conducted an elimination test for up to 5 days. The results showed that the gut retention time (GRT) of MPs is longer in FW than in SW, indicating that MP elimination occurs more quickly in SW than in FW. However, despite shorter GRTs, higher numbers of MPs tended to be retained for a longer period in SW larvae than FW larvae. Subsequently, using a fluorescent marker, gastrointestinal fluid was found to move more rapidly in the SW group. This finding indicates that water drinking accelerates gastrointestinal fluid movement, which moves MPs through the gut in SW larvae. Beside the difference in physiological conditions, MP elimination was faster when food was available, suggesting that feeding also affects MP elimination in fish. Internal factors such as body size and intestine length were also examined, but indicated no significant difference between SW and FW. Therefore, osmoregulation and feeding both influence MP elimination in fish.

## 1. Introduction

Microplastics (MPs), plastic particles smaller than 5 mm, are one of the biggest environmental issues in recent decades (Law and Thompson, 2014). MPs have been detected in both seawater (SW) and freshwater (FW) (Abimbola et al., 2024; Tang et al., 2021). Detection of MPs in aquatic animals has been reported frequently (Ali et al., 2024; Elizalde-Velázquez and Gómez-Oliván, 2021). As some of the most abundant organisms in aquatic ecosystems, fish face serious problems with MP contamination. Ingestion of MPs by fish in natural environments has been extensively reported. MPs accumulate mostly in the gastrointestinal tract and gills of fishes, both marine and FW (Jaafar et al., 2021; Khan et al., 2023; Lam et al., 2022; Lopes et al., 2023).

Fish maintain physiological homeostasis differently in SW and FW environments. Due to osmotic pressure differences, marine fish tend to lose water from their tissues. To compensate for this water loss, marine fish constantly drink water from the surrounding. In contrast, FW fish rarely drink water because they absorb water continually (Evans, 1968; Grosell, 2013). This difference in water drinking behavior for osmoregulation potentially affects MP ingestion in SW and FW fishes. Our previous study reported that marine fishes are more likely to ingest MPs than FW fishes, due to water drinking for osmoregulation (Pratiwi et al., 2023b).

Ingestion of MPs leads to ecotoxicological impacts on fish. However, to better understand MP effects in fish, it is also important to evaluate retention and elimination from the body. Egestion is the process of expelling undigested waste material from the body by excretion or defecation in organisms, thereby reducing adverse effects of pollutants (Ribeiro et al., 2019). Recently, numerous studies have reported MP egestion in various fishes (Cong et al., 2019; Ohkubo et al., 2020; Sun et al., 2023); however, studies focusing on egestion are still limited compared to those concerning ingestion. Therefore, the effect of water salinity conditions, i.e., SW versus FW on MP egestion remains unclear. Also, most studies about MP egestion in fish have been limited to research using MP particles >250 µm; hence, little is known regarding egestion of smaller MPs, although previous studies have reported that factors such as size, shape, and exposure affect MP retention and elimination from intestines of fish (Jabeen et al., 2018; Roch et al., 2021; Santana et al., 2021).

The correlation between digestive system condition and MP retention is an open topic. Previous studies have conducted MP exposure tests by feeding fish with MP-laced food pellets. Results showed that MP retention was not significantly different from those of digesta or food (Grigorakis et al., 2017; Spanjer et al., 2020). However, these studies were limited to experiments using MPs in the size range of 50 - 500 µm. Thus, the relation between retention times of smaller MPs and food availability in the GITs of fish during elimination is also still unclear. Therefore, it is crucial to clarify the basic mechanism of how fish eliminate MPs in SW and FW environments.

In this study, a euryhaline fish, Javanese medaka (*Oryzias javanicus*), was used as a model fish. This species inhabits both SW and FW (Inoue and Takei, 2003; Takehana et al., 2020), and is useful for comparing egestion of MPs in both environments. We exposed 21-day-post-hatching (dph) larvae of *O. javanicus* in SW and FW to 1-µm fluorescent polystyrene (PS) particles for 24 h and investigated egestion of these particles in SW and FW. Beside the difference in salinity, the influence of food was also observed. By investigating the elimination process, kinetic properties of MPs, e.g., gut retention time, elimination time, and elimination patterns were revealed, and the effects of SW, FW, and feeding status were explored. Several factors such as morphological differences and gastrointestinal fluid movement, which may influence MP elimination, were also examined in this study. Moreover, techniques for observing and identifying intestinal fluid movement in larval fish were established in this study. Finally, our study suggests that MP egestion in SW and FW differs due to differences in physiological conditions for fish osmoregulation in both environments.

## 2. Material and Methods

### 2.1. Experimental fish and husbandry

Javanese medaka (*Oryzias javanicus*) strain (RS831) from Penang, Malaysia, were acquired from the National Bioresource Project (NBRP) MEDAKA at the National Institute for Basic Biology, Okazaki, Japan. These fish were maintained in an aquarium system (Iwaki Co., Tokyo, Japan) with recirculating natural seawater. Salinity of seawater for fish husbandry was maintained at ∼31 ppt. Adult and larval fish were fed with brine shrimp larvae (*Artemia nauplii*) (INVE Aquaculture, Belgium) twice a day. The photoperiod was set to 14/10 h day/night at 26℃. Under these circumstances, adult fish spawn every morning. All experimental procedures were approved by the Animal Ethics Committee of the Atmosphere and Ocean Research Institute of the University of Tokyo (permission no. P18-13) in compliance with ARRIVE guidelines.

### 2.2. Larval fish acclimation at different salinities

Larvae of *O. javanicus* (21 days post-hatching; dph) were used in all MP exposure and elimination experiments. Fertilized eggs of *O. javanicus* were obtained by mass mating in 31-ppt SW. Eggs were incubated until hatching in petri dishes containing methylene blue in 31-ppt SW. Acclimation of larvae to SW and FW (aged tap water, 0 ppt) was conducted as described in Supplementary Fig. S1 of Pratiwi et al. (2023b). To remove all gut contents, larvae of both groups were fasted for 24 h before the experiment.

### 2.3. Fluorescent microplastic

Polystyrene MPs (PS-MPs; diameter 1 µm with a coefficient of variation = 2%) labeled with a yellow-green fluorescent dye (excitation 441 nm and emission 485 nm) were obtained from Polyscience, Inc., Warrington, PA, USA (Catalog No. 17154-10) (Fig. 1a). PS microspheres (density: 1.05 g/cm^3^) were provided as a suspension in water and were easily dispersed without aggregation. The PS-MP stock solution was stored at 4°C. Microsphere properties, e.g., fluorescence intensity, diameter, perimeter, were measured using an all-in-one fluorescence microscope (BZ-X800, KEYENCE, Osaka, Japan) equipped with analyzer software (BZ-H4C-Hybrid Cell Count v.1.1.1.8, KEYENCE). Results are shown in Fig. 1b-d.

**Fig. 1.**
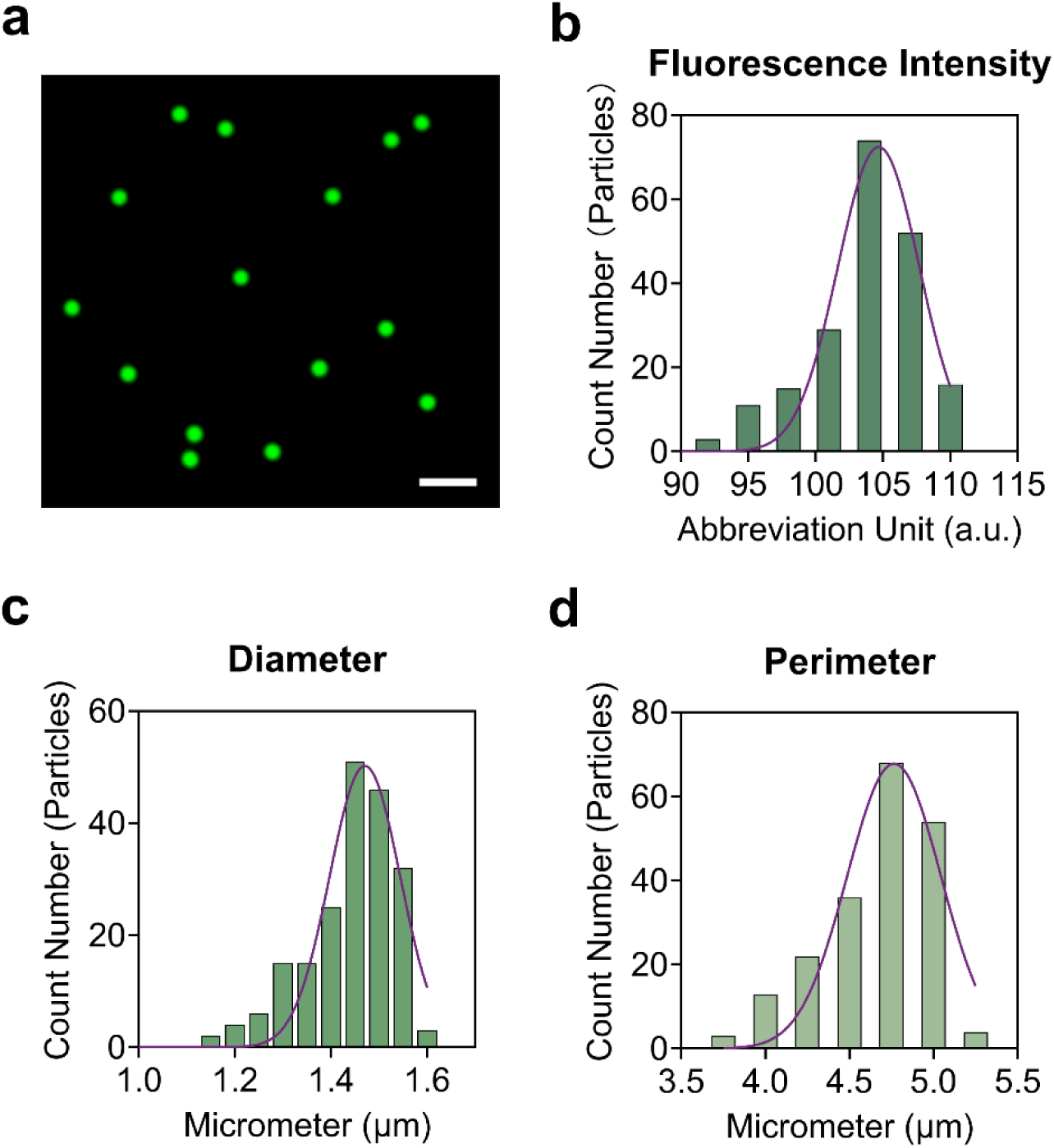
Characteristics of polystyrene fluorescent microspheres (diameter 1.0 µm) used in this study. (a) Microspheres under a fluorescence microscope, (b) Fluorescence Intensity, (c) Diameter, (d) Perimeter. Microplastic characteristics were observed under a fluorescence microscope (KEYENCE) using analyzer software (KEYENCE). Total observed particle numbers n= 199. The scale bar indicates 5.0 µm.

### 2.4. Microplastic exposure test

FW and SW were filtered before use for experiments using a 0.22 µm Millipore sterile vacuum filtration system (Sigma-Aldrich, Germany) to avoid contamination with other particles. 300 mL of filtered SW and FW were poured into 500-mL glass beakers. Exposure solutions were prepared by adding MPs to each beaker to the final concentrations. MP solutions were stirred gently using metal spatulas. Larvae of SW and FW groups (6 larvae per beaker) were carefully transferred into each MP solution using a 2-mL pipette. Each beaker was aerated using an aeration stone to avoid microbead aggregation and to maintain a uniform particle distribution during the exposure test. During exposure tests, medaka larvae in SW and FW (n=66) were exposed to 0.25 mg/L (10^8^ particles/L) of 1-μm PS-MPs for 24 h (Supplementary Fig. S1). This concentration was selected because it was within the range of MP concentrations reported from marine and FW environments (Goldstein et al., 2012; Nunes et al., 2023; Phuong et al., 2022). Larvae for negative controls (n=6) were prepared in both FW and SW without MPs and collected after 24 h. Water temperature and salinity were monitored during exposure tests using a digital salinity meter (Hanna Instruments Inc., HI98319). Densities of FW and SW used in this study were derived from temperature and salinity data (Supplementary Table S1 and S2) using calculation reported previously (UNESCO, 1981). Subsequently, dynamic viscosity of FW and SW (Supplementary Table S1 and S2) were also calculated, as described in Kestin et al. (1978).

### 2.5. Microplastic elimination test

After MP exposure, body surfaces of larvae were wiped carefully using a paper towel to remove attached MPs. Larvae from both groups were transferred to glass beakers containing 200 mL MP-free SW or FW. A total of 12 larvae were placed in a beaker for the short-term elimination test while 54 larvae were placed in another beaker for the long-term elimination test. During elimination tests, food was not distributed to larvae in the no-food groups. To prevent reingestion of excreted MPs by larvae, all glass beakers were equipped with a stainless net and each larva was placed in it during the elimination test (Supplementary Fig. S1). In the short-term elimination test, to examine the number of particles excreted by larvae at each time point, water was sampled 0, 3, 6, 9, 12, and 24 h after transfer. Additionally, all larvae from both groups were also collected at 24 h to count the remaining MPs. In the long-term elimination test, larvae exposed to MPs were reared in MP-free SW or FW for up to 5 days. Six larvae per day were sampled on days 0, 1, 3, and 5. During the long-term elimination period, water from each beaker was sampled every day before water replacement.

Feeding status may also affect MP elimination in the larval gut. To examine this possibility, 66 larvae were prepared for a similar exposure test followed by 24-h short-term elimination test (n= 12 larvae) and a 5-day long-term elimination test (n= 54 larvae). Larvae in SW and FW were fed with artemia larvae for 30 min before the elimination test. The elimination test and sampling procedure were generally conducted as in the non-feeding group experiment. Room temperature during the elimination test was maintained at 26℃. Water conditions during experiments were monitored by placing an identical glass beaker filled with 200 mL SW or FW. Water temperature and salinity were measured and then densities and viscosities of water at each time point during the short and long-term elimination phases were calculated using the methods described above (Supplementary Table S1 and S2).

### 2.6. Microplastic extraction from larvae and water samples

All larvae collected during exposure and elimination tests were sacrificed on ice. After sacrifice, each larva was put in a 1.5-mL Eppendorf tube and homogenized in 10% KOH solution using homogenizing pestles. Subsequently, solutions containing homogenized tissues were mixed well by pipetting, and then incubated at 42°C for 3 days to degrade tissues. To extract 1-μm PS-MPs, digested tissue solutions were sonicated and filtered using 0.45 µm IsoporeTM polycarbonate membrane filters with vacuum filtration. In addition to larval samples, 200-mL water samples were sonicated to separate MPs from feces for MP extraction during the short-term elimination phase. Sonicated water samples were also filtered using 0.45 µm IsoporeTM polycarbonate membrane filters with vacuum filtration. Meanwhile, 200 mL of water samples containing fecal pellets and MPs from the long-term elimination phase were vacuum-filtered without sonication to observe MP accumulation in fecal pellets. All membrane filters were dried overnight and then set on glass slides with coverslips for observation and particle counting.

### 2.7. Detection and Automatic counting of MPs

Detection and counting of MPs on filter membranes were performed according to a previous report (Pratiwi et al., 2023b) using an all-in-one fluorescence microscope (KEYENCE) equipped with a GFP filter (excitation 470/40 nm, emission 525/50 nm, dichroic 495 nm). Numbers of eliminated MPs at each sampling point during short and long-term elimination were counted automatically. In addition, total ingested particles were calculated by summing numbers of eliminated MPs at 3, 6, 9, 12, 24 h with the number of MPs remaining in larval bodies at 24 h. Moreover, MP distribution in larval bodies after the exposure test was confirmed by transparentizing larval tissue using a procedure reported previously (Pratiwi et al., 2023b).

### 2.8. Estimation of gut retention time

Gut retention time (GRT) denotes the length of time required to eliminate digesta or other materials from the gut (Stevens and Hume, 2004). In this study, GRTs of MPs were estimated as GRT50, GRT90, and GRT95, the times to eliminate 50%, 90%, and 95% of MPs from larval bodies, respectively. According to guidelines for bioaccumulation tests in fish (OECD, 2012), GRTs were calculated with the following equation:

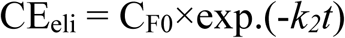

where CE_eli_is the MP concentration in the fish at hour *t* of the elimination experiment (particles/g wet weight), and C_F0_ is the MP concentration in the fish at the beginning of the elimination experiment. In this equation, *k_2_* is the elimination rate constant, which represents the fraction of MPs eliminated per unit time in each species.

In this study, calculation of GRTs was based on MP numbers counted at each sampling point during the short-term elimination phase. Moreover, the linear regression model was used to predict the relationship between MP concentration and excretion time in this study.

### 2.9. Live imaging observations of water drinking and gastrointestinal fluid movement in larval fish

In teleost fishes, melanin develops in the abdomen commencing in the early larval stage and becomes more intense 3 weeks post-hatching (McMenamin et al., 2014; Pratiwi et al., 2023a), which restricts live imaging observations in the gastrointestinal area. However, transparentization is not applicable to tracking imbibed water because fluorescent dye is washed out during treatments. Therefore, instead of 21 dph larvae, 4 dph larvae were used to observe water drinking and gastrointestinal fluid movement in the SW and FW groups. Before the experiment, some hatched larvae (n=6) were reared in SW and others (n=6) were acclimated in FW for 4 days. Larvae of SW and FW groups were immersed in the two media containing 1-µM fluorescein isothiocyanate (FITC)-labelled dextran (Average Molecular Weight 70,000, Sigma, 46945), which acts as a marker for drinking. After 1 h of immersion, larvae were removed and rinsed with FW or SW for 5 min to remove excess FITC-dextran. Each larva was observed at 0, 2, 3, and 24 h after immersion by moving it to a 1-cm hole in a 2% agarose gel made in a glass-based dish. During observations, the signal of FITC-dextran in larval intestines was examined from a ventral orientation focusing on the abdomen to the cloacal and was imaged in 3D using a fluorescence microscope. Larvae were returned to a petri dish containing SW/FW as quickly as possible after observation. After 24 h, larvae were collected and stored in 10% Formalin at 4℃ for body measurements.

Water influx in *O. javanicus* larvae in SW and FW was estimated by analyzing 3D images of FITC-dextran signals observed at 0 hour using 3D measurement KEYENCE analyzer software. These images of FITC-dextran signals in larval GITs were then imported to Fiji (ImageJ). Subsequently, fluorescent signals detected from all areas of GITs were analyzed using plot profiles. The average value of fluorescence intensity relative to distance (pixels) in all images was calculated. Afterward, the location of each fluorescent signal was identified by calibrating the distance on the plot to the somite number on the image. Furthermore, fluorescence gradients from all samples were combined in GraphPad Prism 10 using a heatmap, and water movement patterns were compared between SW and FW groups.

### 2.10. Measurement of larval total body length and gastrointestinal tract length

Gastrointestinal tracts (GITs) were dissected from 21 dph larvae of SW and FW groups after anesthetization on ice. The whole body and GITs of both groups (n= 20) were fixed in 10% formalin for 3 days. Before measurements, 10% formalin was discarded and replaced with PBS solution for sample washing. Lengths of whole bodies and GITs were observed on a glass-bottomed dish filled with glycerol and then measured from rostral to caudal axis using KEYENCE analyzer software.

### 2.11. Statistical analysis

All counting and measurement data were summarized in Microsoft Office 2010 and then statistically analyzed in GraphPad Prism 10 (Version 10.2.3., 2024). For GRT calculation and MP accumulation, significant differences between SW, FW, feeding and non-feeding groups were analyzed using one-way ANOVA followed by Tukey’s Honestly Significant Difference Post-Hoc test. For comparative analysis of water influx, total body length, and intestine length, significant differences between SW and FW groups were analyzed using Student’s t-test. The level of statistical significance was set at P < 0.05 in all analyses.

## 3. Results

3.1. Microplastic elimination timing and pattern in *O. javanicus* larvae during a short-term elimination test

Microplastic elimination was observed for 24 h in SW/FW larvae without food. Overall excretion patterns differed among 12 larval fish in both groups, although all larvae gradually eliminated MPs for up to 24 h at all sampling time points (Supplementary Fig. S2 and S3). In the SW group, most larvae started to excrete MPs as early as 3 h after transfer to MP-free SW. High numbers of MPs were excreted within 9 h after transfer (Fig. 2a). In the FW group, most larvae also began to eliminate MPs at 3 h. However, most FW larvae excreted a high number of MPs around 12 h after the transfer, later than the SW group (Fig. 2b).

**Fig. 2.**
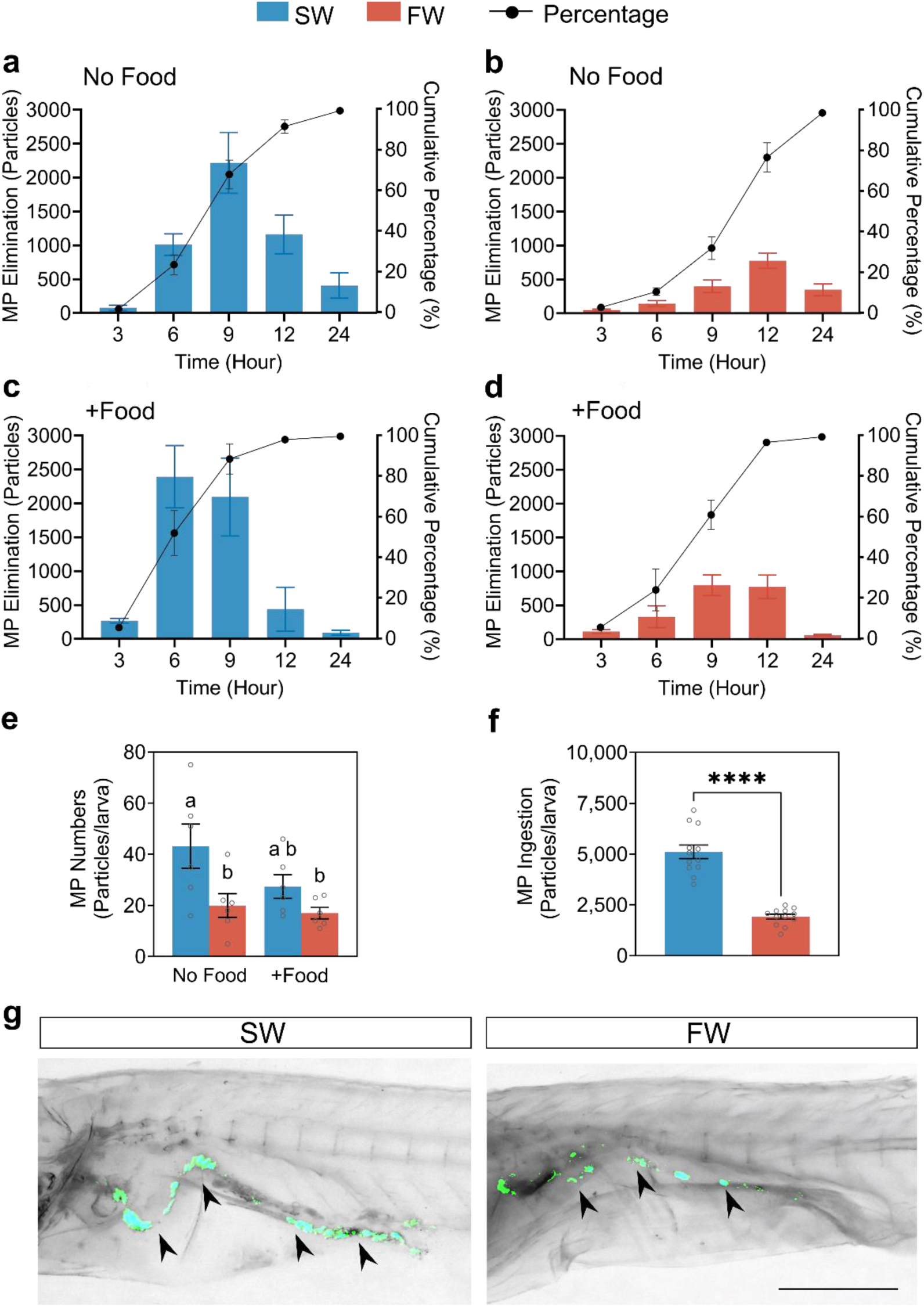
Microplastic ingestion and elimination in *Oryzias javanicus* larvae. Microplastic elimination pattern and timing in (a) the Seawater (SW)-non-feeding group, (b) Freshwater (FW)-non-feeding group, (c) SW-feeding group, (d) FW-feeding group. Each bar shows the mean of particle numbers of eliminated MPs ± standard error of the mean (SEM) at each time point, whereas the line represents the mean cumulative percentage of MP elimination ± SEM. Number of fish observed n=6. (e) Accumulation of microplastic in larvae 24 h after the elimination test. Each bar shows the mean number of MPs remaining in the larval body ± SEM. Different letters above the bars indicate significant differences, which were analyzed using a one-way analysis of variance (ANOVA) (P < 0.05) followed by Tukey’s Honestly Significant Difference post-hoc test. (f) The number of ingested MPs after a 24-h exposure test. Each bar shows the mean value ± SEM. Differences between groups were analyzed for significance using Student’s t-test (ns, non-significant; P > 0.05), with P<0.0001 shown as ****. (g) Distribution pattern of ingested MPs in gastrointestinal tracts of SW- and FW-*O. javanicus* larvae. Arrowheads show MP particles in the intestine. The scale bar indicates 0.5 mm.

All larvae fed with artemia before the elimination test also initially eliminated MPs 3 h after the transfer (Supplementary Fig. S4 and S5). High numbers of MPs were excreted relatively faster than in the non-feeding groups, 6-9 h in the SW group and 9-12 h in the FW group (Fig. 2c, d). Moreover, MPs were still found in all larvae with food after the 24-h elimination test. The mean concentration of remaining MPs was quite similar in larvae without food and larvae with food, but significantly higher in SW larvae (Fig. 2e). Numbers of ingested MPs in both media were calculated from total eliminated MPs and MPs remaining in larval bodies. SW larvae ingested significantly higher numbers of MPs than FW larvae (Fig. 2f, g).

### 3.2. Estimation of gut retention time (GRT50, GRT90, GRT95) during a short term-elimination test

Gut retention times (GRTs) were estimated by linear regression, followed by a calculation using an elimination equation used for individual larvae (Supplementary Fig. S6-S9). In comparisons between the SW and FW groups, GRT50 and GRT90 were significantly longer in the FW group than the SW group (Fig. 3a, b). The GRT95 in FW groups was also longer than in SW groups, although not statistically significant (Fig. 3c). Subsequently, a comparison between feeding status showed that all GRTs were longer in non-feeding groups than feeding groups regardless of water type. Overall, MP retention was estimated to be longer in FW than in SW. Also, these results imply a tendency of larvae with food to excrete MPs more quickly than larvae without food in their GITs (Fig. 3a-c). Beside GRTs, the elimination rate constant k_2_ of 1 µm PS-MPs used in this study was also estimated and compared among larvae. In non-feeding groups, the value of k_2_ was 5.2 day^-1^ in SW larvae and 4.5 day^-1^ in FW larvae. Meanwhile in feeding groups, the value of k_2_ was 5.5 day^-1^ in SW larvae and 5.4 day^-1^ in FW larvae. Overall, k_2_ was larger in SW larvae, especially in the feeding groups (Supplementary Table S3). As with GRT values, this result indicates faster elimination of MPs in SW and feeding groups.

**Fig. 3.**
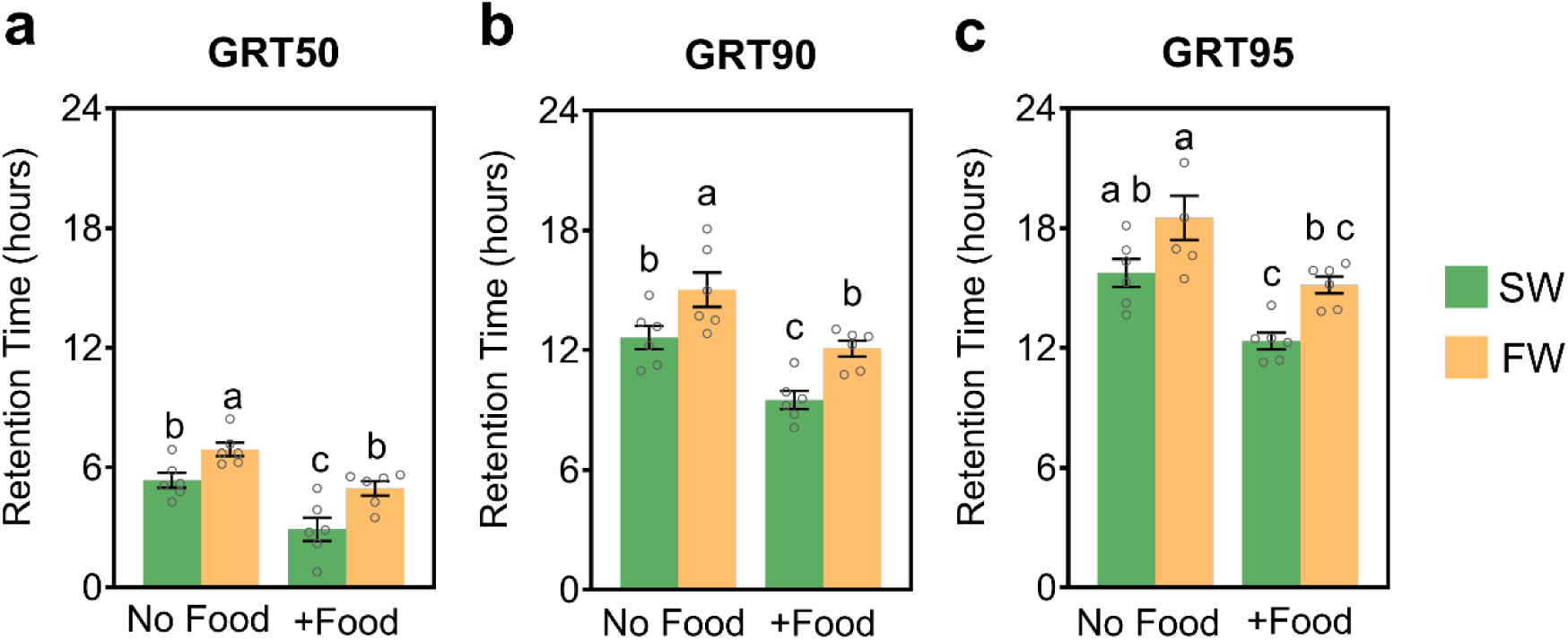
Comparison of gut retention time (GRT) among all groups. (a) GRT50, (b) GRT90, (c) GRT95. Each bar shows the mean number of MPs remaining in the larval body ± standard error of the mean (SEM). Different letters above bars indicate significant differences, analyzed using one-way analysis of variance (ANOVA) (P < 0.05) followed by Tukey’s Honestly Significant Difference post-hoc test.

### 3.3. MP detection in larval bodies and fecal pellets during the long-term elimination test

Numbers of MPs accumulated in the larval body and excreted via fecal pellets during long-term elimination were also observed. MP numbers in the larval body were higher in SW than FW in all sampling days, especially in non-feeding groups (Fig. 4a). Also, all larvae in feeding and non-feeding groups showed that MPs could not be fully eliminated until Day 5. However, larvae with higher MP numbers were predominantly non-feeding SW larvae (Fig. 4a). In addition, observation of fecal pellets showed that most MPs were excreted with fecal pellets on Day 1 (Fig. 4b). Overall, numbers of MPs decreased after Day 1, but still could be detected until Day 5 in fecal pellets of SW larvae of both feeding groups (Fig. 4b).

**Fig. 4.**
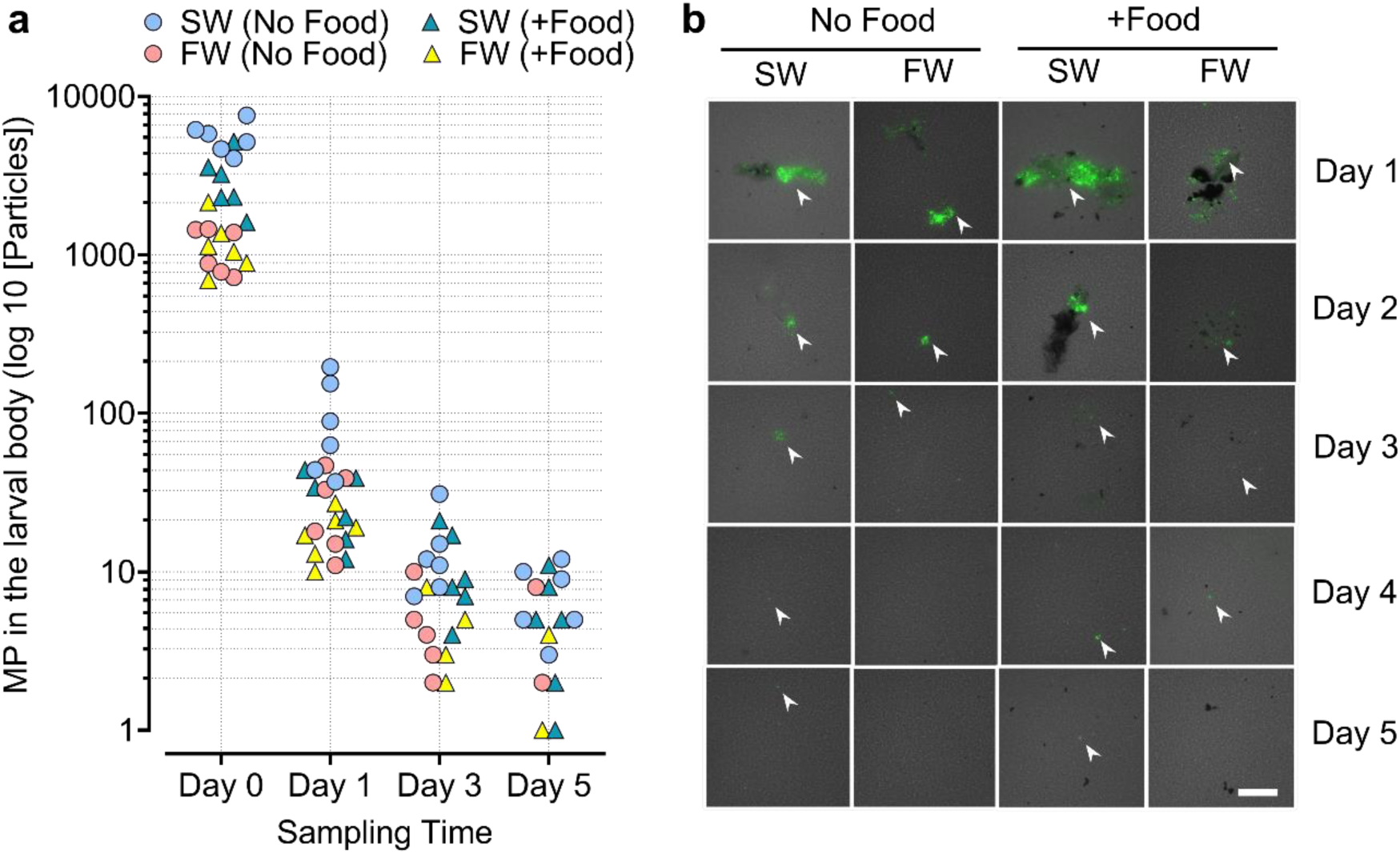
Microplastic accumulation in larvae and fecal pellets during the long-term elimination test. (a) Microplastic accumulation in the larval body during long-term elimination tests. Each dot represents numbers of MPs accumulated in each larval body. (b) Microplastic accumulation in fecal pellets during long-term elimination tests. White arrows represent signals of MPs in fecal pellets. The scale bar indicates 100 µm.

### 3.4. Comparison of gastrointestinal fluid movement in SW and FW larvae

Gastrointestinal fluid movement was observed by exposing larvae to SW/FW containing FITC-dextran to visualize water drinking (Fig. 5a). Green fluorescent signals of FITC-dextran in larval GITs were observed at 0, 2, 3, and 24 h after exposure. Positions of fluorescent signals were located using somites in larval bodies as a scale. Then they were compared between SW and FW larvae. In the SW group, FITC-dextran signals were detected between somites 2 and 6 at 0 h. Signals were shifted posteriorly and reached somite 8 at 2 h and reached the cloacal area (somite 9) within 3 h (Fig. 5b, c). Meanwhile, in the FW group, at 0 hour, FITC-dextran signals were detected between somites 1 and 5. Stronger signals were detected between somite 2 and somite 5 in FW larvae at 2 h. However, no signal was detected around the cloacal area at 3 h, yet signal was found between somites 2 and 6 (Fig. 5b, c). Additionally, after 24 h, highly visible FITC-signals were observed at the cloacal area in SW larvae. However, compared to FITC-dextran signals in SW larvae, which completely disappeared after 24 h, some FITC-dextran signals remained in the anterior part of FW larvae (Fig. 5b, c). Furthermore, in this study, water influx of *O. javanicus* larvae was also estimated from fluorescence signals at 0 h. Larvae in SW ingested water 4-5 times more than those in FW (Fig. 6a).

**Fig. 5.**
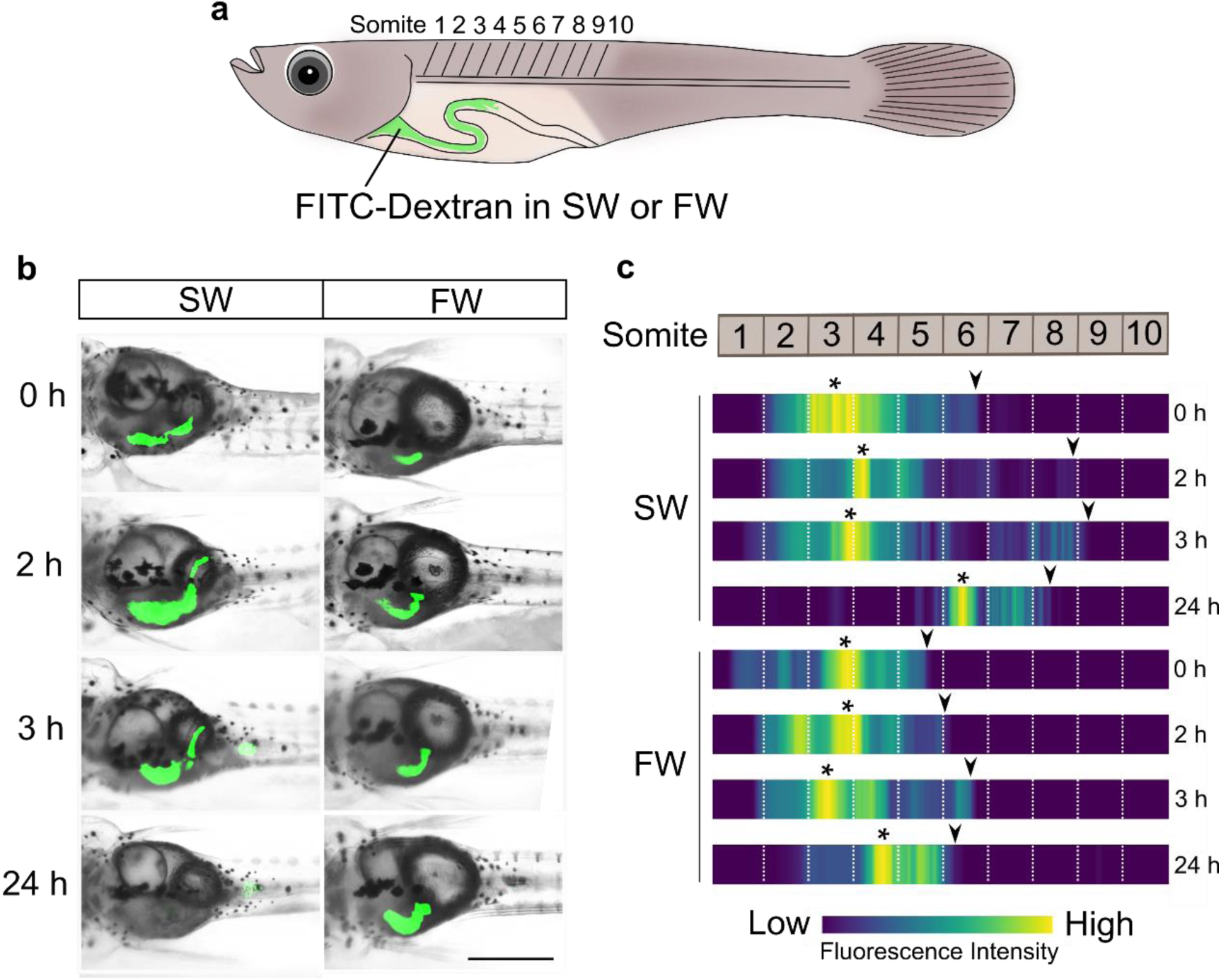
Observation of gastrointestinal fluid movement in *Oryzias javanicus* larvae for up to 24 h after 1 h of immersion in FITC-dextran. (a) Positions of the gastrointestinal tract of *O. javanicus* larvae relative to somite numbers. (b) Distribution of water containing the fluorescent signal of FITC-dextran in the gastrointestinal tract of larvae in the SW and FW groups. (c) Fluorescence gradient of FITC-dextran signals in SW and FW larvae. Asterisks represents signal peaks, and arrowheads represents the most posterior part of the signal in gastrointestinal tract. The scale bar indicates 0.5 mm.

**Fig. 6.**
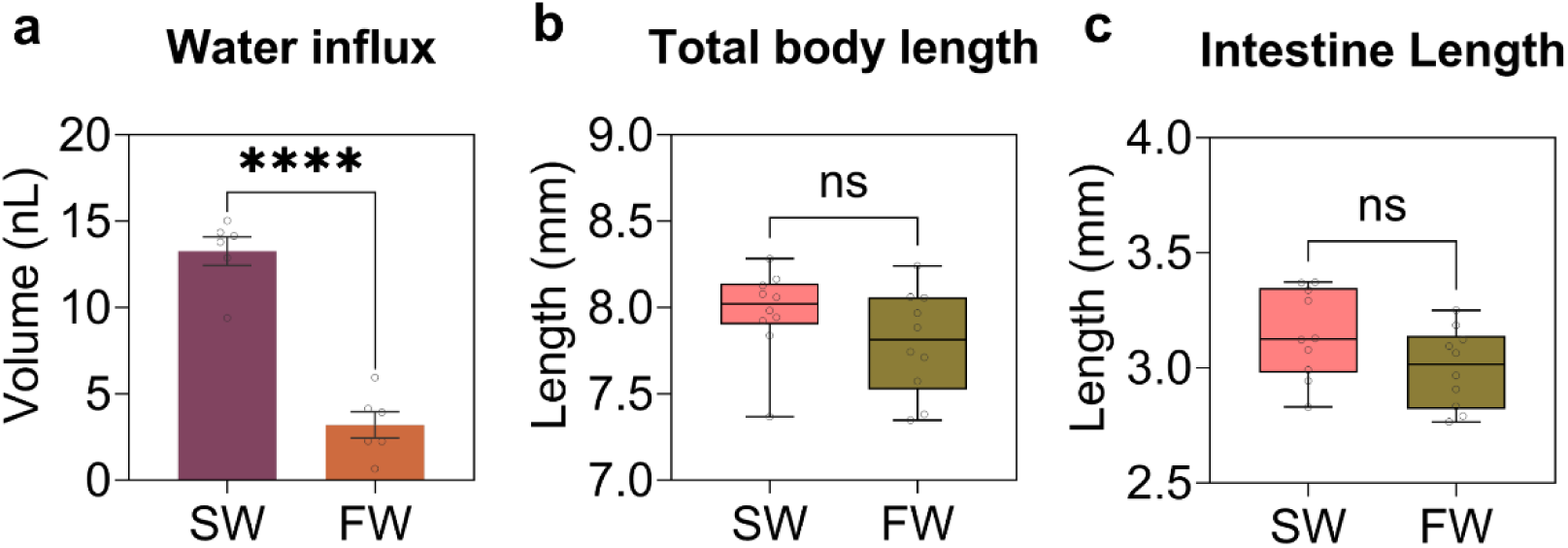
Factors that may affect MP elimination in larval fish. Comparison of (a) Water Influx, (b) Total body length, (c) and Gastrointestinal tract lengths of SW and FW larvae. Each bar and box plot shows the mean ± the standard error of the mean (SEM). Differences between groups were analyzed for significance with Student’s t-test (ns, non-significant; P > 0.05), with P<0.0001 shown as ****.

## 4. Discussion

### 4.1. Osmoregulation influences gastrointestinal fluid movement in fish

Using *Oryzias javanicus* larvae as a model fish, this study clearly demonstrated that fish in SW and FW eliminate MPs at different rates. Our findings indicated that gut retention times of MPs are longer in FW fish; thus, MP elimination tends to occur more quickly in SW fish than in FW fish (Fig. 3a-c). This suggests that osmoregulatory differences also affect MP elimination processes in SW and FW larvae, in addition to effects on MP ingestion previously reported (Pratiwi et al. 2023b).

To compensate for water loss, SW fish continuously drink water in high salinity environments. Most MPs ingested in SW went to the intestine because the esophagus of medaka fish is short, and the contribution of the stomach to ion and water regulation is generally minor (Takei, 2021). In the intestine, ingested SW is desalinized and then absorbed. Therefore, luminal water movement is active in intestines of SW fish, especially in the anterior intestine, although the amount of water decreases in the posterior intestine (Grosell et al., 2010). In contrast, FW fish drink less water because water easily enters their tissues by osmosis from body surfaces, including the gill epithelium. Absorbed water is excreted by the kidney as urine. The GIT is not involved in this process. Therefore, smaller amounts of water pass through the GIT in FW fish (Grosell et al., 2010).

Osmoregulatory differences affect gastrointestinal tract functions in fish. A study using euryhaline teleost rainbow trout (*Oncorhynchus mykiss*) showed that intestinal contractile activity dramatically increased upon seawater exposure (Brijs et al., 2017). Moreover, salinity is also reported to affect digestion in milkfish (*Chanos chanos*), in which intestinal passage time is significantly faster in SW than FW fish (Ferraris et al., 1986). These reports suggest that the increase in water-drinking activity influences the function of the gastrointestinal tract. In this study, using FITC-dextran, observations of gastrointestinal fluid movement in *O. javanicus* larvae (4 dph) showed that gastrointestinal fluid moved more rapidly in the SW group and reached the cloacal area within 3 h after immersion (Fig. 5b, c). SW fish absorb as much water as possible to adjust their osmolality; thus, intestinal fluid moves more rapidly in SW fish than in FW fish, which may influence retention and elimination of MPs.

### 4.2. Food in the gastrointestinal tract accelerates MP elimination

Compared to ingestion, how fish eliminate MPs is more complicated and has not been understood. MP properties, such as size, shape, concentration, and exposure duration, have been reported as factors that affect MP elimination in fish (Abarghouei et al., 2021; Mak et al., 2019; Roch et al., 2021; Zhang et al., 2021). Also, internal factors such as gut morphology reportedly influence MP elimination in fish (Ohkubo et al., 2022). In addition to these factors, we considered osmoregulation differences and feeding status, which might also contribute to MP elimination in fish.

Thus far, the correlation between MP retention time and food availability in the GIT of fish is still unclear. Information is restricted to studies that were conducted using food-size MPs in the presence of food in GITs of fish. For example, studies using goldfish (*Carassius auratus*) and Japanese anchovy (*Engraulis japonicus*) reported that retention of MPs (diameter 250-300 µm) was generally similar to retention of food or digesta in GITs of control fishes (Grigorakis et al., 2017; Ohkubo et al., 2022). Calculation of GRTs in this study estimated that MP retention is likely longer in non-feeding larvae than in feeding larvae (Fig. 3a-c). Previously, Liu et al. (2021) demonstrated that MPs with a diameter of 2 µm have a longer retention time in intestines of juvenile Japanese medaka, compared to MPs 20 and 200 µm in diameter. The diameter of MPs used in the present study was 1 µm, which is similar to 2 µm, but considerably smaller than 20 or 200 µm. This indicates that very small MPs may have different kinetic behaviors than food-size MPs. Thus, food availability in the intestine can influence retention of very small MPs in fish.

Food presence was reported to promote contractions of the gut/intestines and peristaltic actions, which accelerate digestion in fish (Olsson and Holmgren, 2001; Smith, 1980). In this study, MP elimination was faster with food in the GIT of both SW and FW larvae. In the non-feeding group, the highest number of excreted MPs was generally detected within 9 h in the SW group and at 12 h in the FW group (Fig. 2a, b). However, high numbers of MPs tended to be eliminated earlier in feeding groups, mostly from 6-9 h in SW and 9-12 h in the FW group (Fig. 2c, d).

Observations of fecal pellets from water samples during the long-term elimination phase showed that excreted MPs were mostly detected in feces rather than free on membrane filters. For instance, an image of fecal pellets from water samples on Day 1 in the SW group indicates that one fecal pellet contained a high abundance of MPs (Fig. 4b). These results imply that fish eliminate most ingested MPs, through fecal excretion not as free particles. Feeding and food presence led to contractions of the stomach and intestine, followed by peristaltic actions, migrational motor complexes, or any other motility pattern (Olsson and Holmgren, 2001; Smith, 1980). Diet contents and food availability in the intestine have also been reported to affect defecation in fish (Rahman et al., 2016; Storebakken et al., 1998). Differences in feeding status can promote peristaltic movement of the GIT and fecal production, which seem to support MP excretion. Therefore, the presence of food in the GIT increases the probability of rapid MP excretion.

### 4.3. MP elimination in various fishes

Fish are one of the most studied aquatic animals in MP research (De Sá et al., 2018). Studies on kinetic properties of MPs during elimination using various fish showed that larger MPs tend to be more quickly excreted by fish (Liu et al., 2021; Ohkubo et al., 2022), whereas the opposite occurs in filter-feeders such as mussels (Kinjo et al., 2019). In this study, elimination of relatively 1-µm MPs showed that *O. javanicus* larvae in all groups could excrete more than 95% of ingested MPs within 24 h, irrespective of salinity or feeding status. Despite very small size, the time required by *O. javanicus* larvae to eliminate more than 95% of 1-µm PS-MPs was relatively similar to findings of previous studies using larger MPs (200∼300 µm) in various fish. For example, Mummichog and Red Seabream reportedly eliminate 95% of ingested MPs within 25 h (Ohkubo et al., 2020). Similarly, a study using both marine and freshwater fishes, such as zebrafish, Japanese medaka, clown anemonefish, and Indian medaka, reported that more than 50% of fishes excreted all ingested MPs within 24 h (Okamoto et al., 2022). These findings imply that MP elimination in fish may be considerably faster than in other animals. In addition, previous research suggested that excretion patterns and timing of MPs in fish is inconsistent and varies widely among individuals of the same species (Okamoto et al., 2022). Similarly, MP excretion patterns in this study also differed among individuals (Supplementary Fig. S2-S5).

Elimination rates of MPs in various fish species have been estimated in several studies, using the elimination rate constant *k_2_* and half-time *T_50_*as parameters. Elimination of 1-µm PS-MPs used in this study were compared among other fish species and particle sizes using these parameters (Supplementary Table S3). In general, comparison of *k_2_* and *T_50_* values between SW and FW species showed that SW fishes tend to have a larger *k_2_* values and a smaller *T_50_*, which indicates more rapid elimination of MPs than in FW species. This result is consistent with findings obtained in this study, which suggest that MP elimination occurs more rapidly in SW environments. Interestingly, comparisons between similar habitats of fish indicates that larger particles likely have larger *k_2_* values; thus, MPs tend to be eliminated more quickly than smaller particles (Supplementary Table S3).

This result is consistent with previously reports using mussels and fish, that MP elimination is size-dependent (Kinjo et al., 2019; Liu et al., 2021). However, comparisons between *Oryzias* fishes in FW showed that *k_2_* in juvenile *O. latipes* was 0.76 day^-1^ (Liu et al., 2021), which is considerably lower than FW-feeding groups of *O. javanicus* larvae estimated in this study, although both studies used relatively similar sized MPs (Supplementary Table S3). Differences in body size, specifically in GIT length, between juvenile and larval fish used for experiments is one possible factor. For reference, the body length of 2-month juvenile *O. latipes* is about 2-3 cm (Iwamatsu, 2004; Kinoshita et al., 2009), whereas the total body length of *O. javanicus* larvae used in this study is 7 - 8.5 mm (Fig. 6b). Accordingly, larval GITs are estimated to be shorter than juvenile GITs.

Thus, MP elimination in this study occurred more rapidly than in an earlier study by Liu et al. (2021). The difference in exposure test duration may also affect the elimination rate of MPs between both *Oryzias* species in the same habitat.

### 4.4. Other factors that influence MP elimination in fish

Beside osmoregulation and feeding status, several factors, such as water temperature and GIT characteristics, have also been reported to influence digestion in fish (Handeland et al., 2008; Kounna et al., 2021; Smith, 1980). In this study, length of the body and the dissected GIT were measured and compared between SW and FW groups. Results (Fig. 6b, c) showed no statistically significant differences. Water conditions such as temperature, salinity, density, and viscosity were also measured and calculated during exposure and elimination tests. Data are available in Supplementary Table S1 and S2, showing that the temperature and salinity of SW and FW used in all elimination tests were relatively constant. Therefore, external factors such as water temperature differences during experiments and morphological differences may not contribute to differences in MP elimination in SW and FW larvae in this study.

In this study, 1-µm MPs were not entirely eliminated from larval bodies (Fig. 2e), suggesting that smaller MPs may persist longer in fish. Consistently, previous studies using 1-or 2-µm MPs also found that smaller MPs tend to remain longer in the bodies of juvenile and larvae *O. latipes* (Liu et al., 2021; Pratiwi et al., 2023c). The intestine possesses villi, increasing the intestinal surface area, so as to enhance absorption (Helander and Fändriks, 2014). This raises the possibility that MPs may become trapped in villi, increasing their residence time. Additionally, although MP elimination tends to occur faster in SW larvae, the number of remaining MPs was also higher in SW larvae than FW larvae (Fig. 2e and 4a). Since SW larvae ingested more MPs than FW larvae during exposure (Fig. 2f), it is convincing that fish that ingested higher numbers of MP are more likely to accumulate MPs in the intestines. Therefore, impacts of MPs are also presumably higher in SW fish than FW fish.

## 5. Conclusions

All findings in the present study indicate osmoregulation as an important factor influencing MP elimination in fish under different salinities. Most ingested MPs were eliminated more rapidly in SW larvae due to the active motility of intestinal fluid in SW environments. Beside physiological factors, food availability in the intestine may also affect MP elimination in fish, i.e., MP elimination occurs more rapidly in fish with food in the intestines. Results of this study are applicable to most teleost species in various osmotic environments because osmoregulatory mechanisms are essentially similar between larvae and adult fish and among teleosts.

## CRediT authorship contribution statement

HMP: Conceptualization, Methodology, Validation, Formal analysis, Investigation, Resources, Writing―original draft, Writing―review & editing, Visualization, TT: Methodology, Validation, Supervision, Writing―review & editing, SR: Resources, Writing―review & editing, KI: Conceptualization, Funding acquisition, Supervision, Writing―review & editing.

## Data availability

All additional data are provided in the Supplementary Data or are available upon request, respectively.

## Declaration of competing interest

The authors declare no competing interests

## Acknowledgement

This work was supported by The University of Tokyo FSI-Nippon Foundation Research Project on Marine Plastics, and JSPS Core-to-core CREPSUM JPJSCCB20200009 to K.I. The authors sincerely thank Mr. Abdul Rauf for helping with fish husbandry.

## Notes

### Competing Interest Statement

The authors have declared no competing interest.

