## Supplementary File 1 for "Osmoregulation affects elimination of microplastics in fish in freshwater and marine environments"

*To whom correspondence should be addressed.

Corresponding Hilda Mardiana Pratiwi

Page 1-15

Supplementary Figure S1-S11

Supplementary Table S1-S3

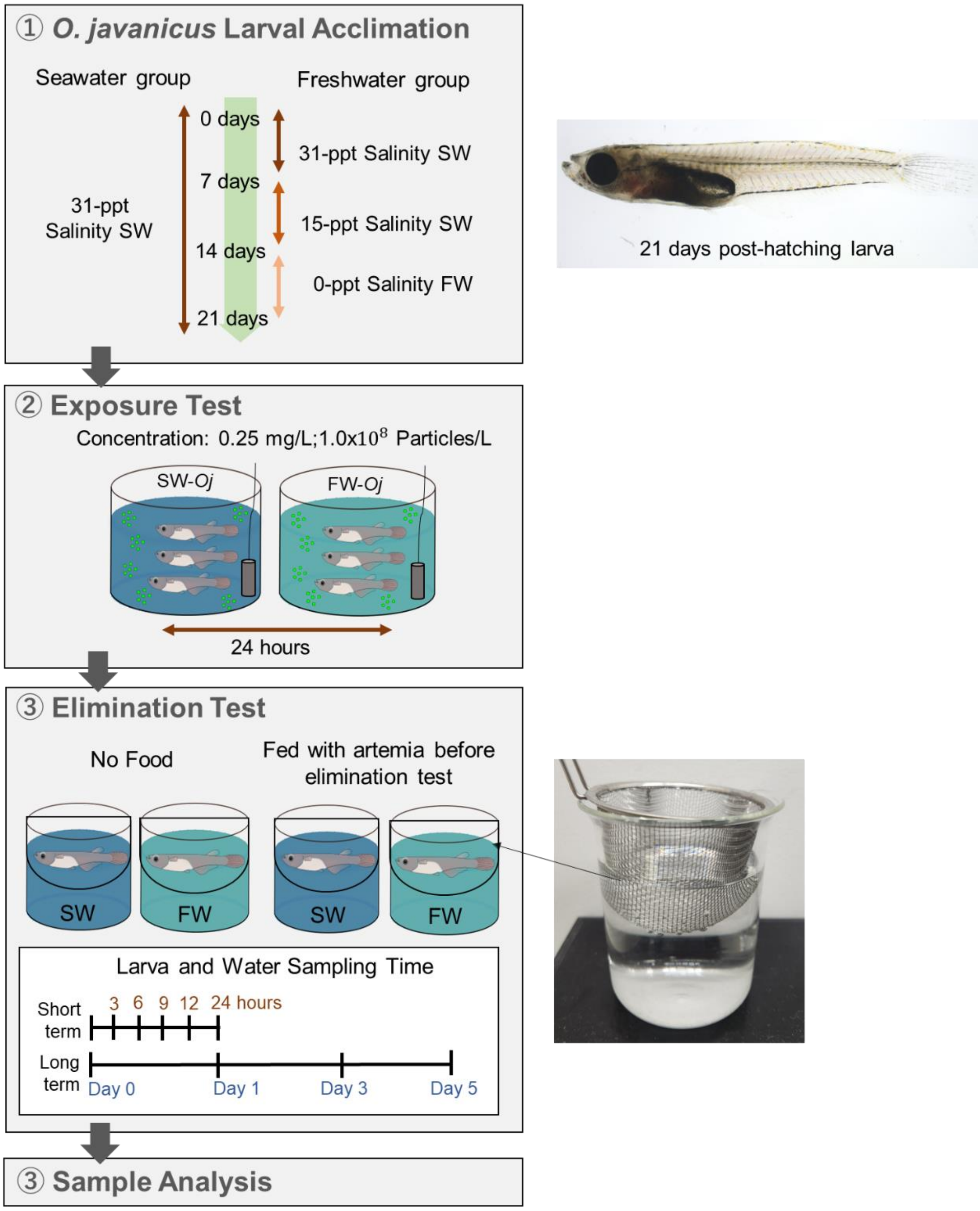


**Supplementary Figure S1. Experimental design of microplastic elimination test involving *O. javanicus* larvae.**

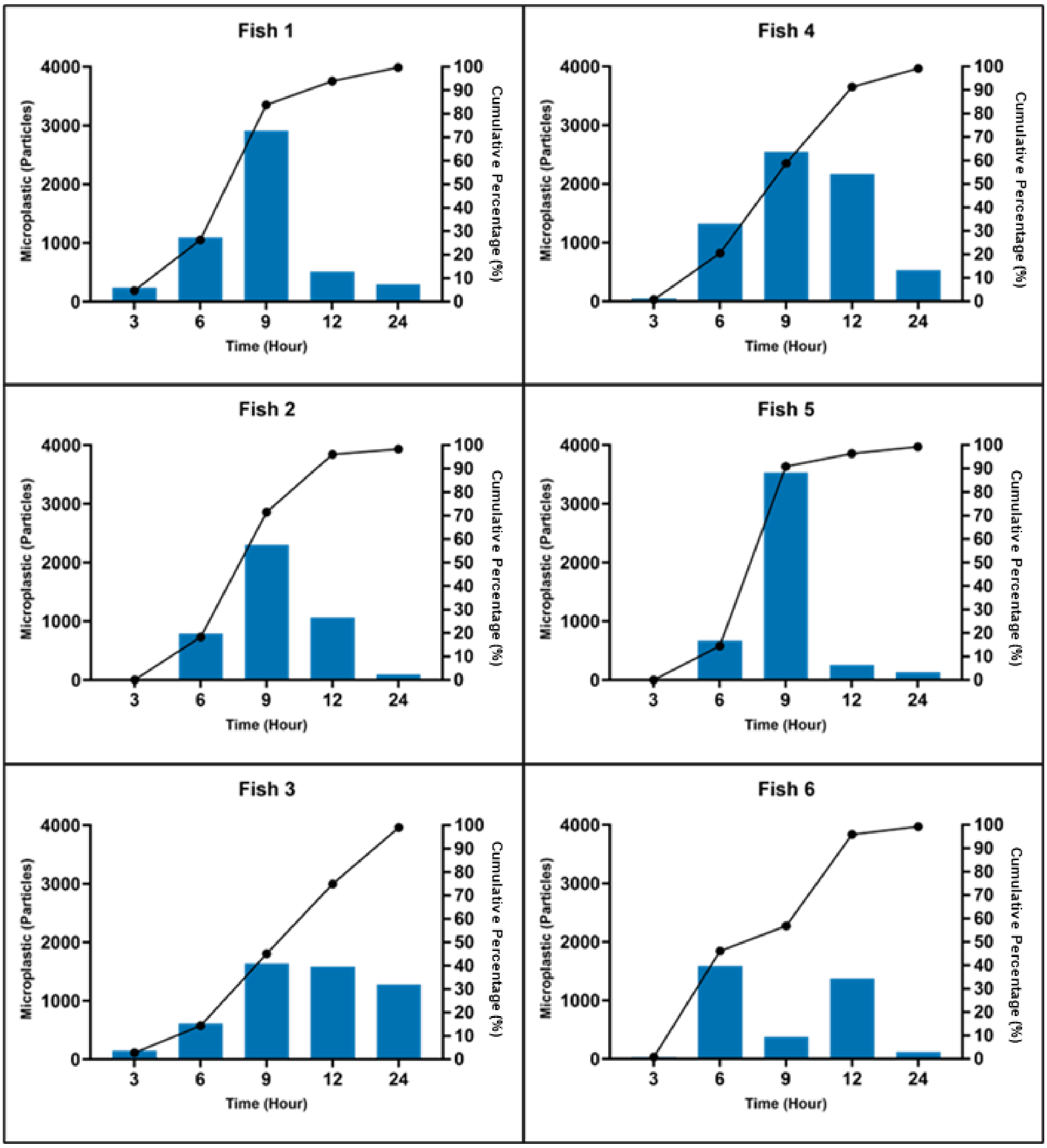


**Supplementary Figure S2. Microplastic elimination and timing in non-feeding *Oryzias javanicus* larvae in seawater.** Each bar shows numbers of eliminated MPs at each time point, whereas the line represents the cumulative percentage of MPs eliminated.

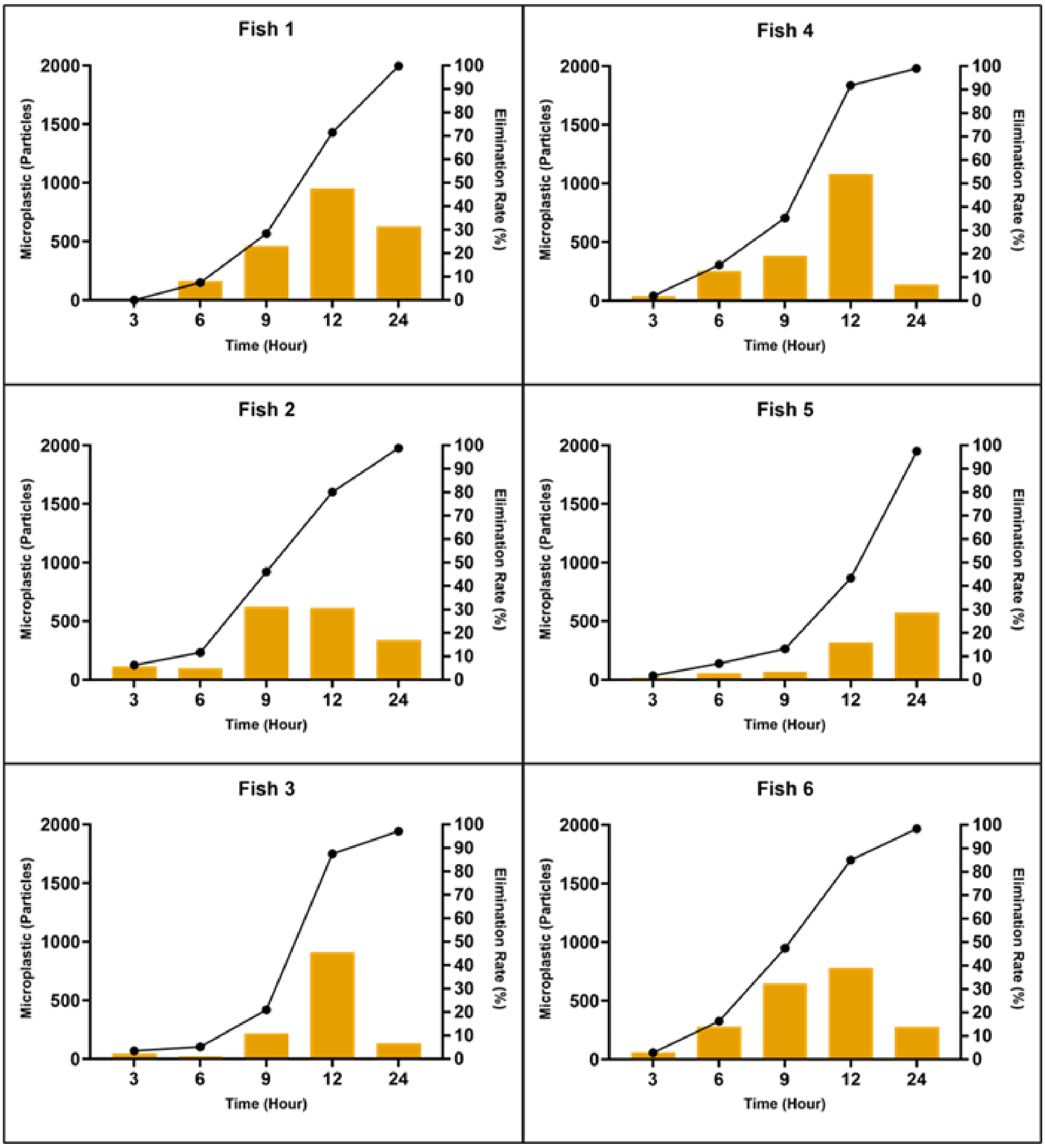


**Supplementary Figure S3. Microplastic elimination and timing in non-feeding *Oryzias javanicus* larvae in freshwater.** Each bar shows numbers of eliminated MPs at each time point, whereas the line represents the cumulative percentage of MPs eliminated.

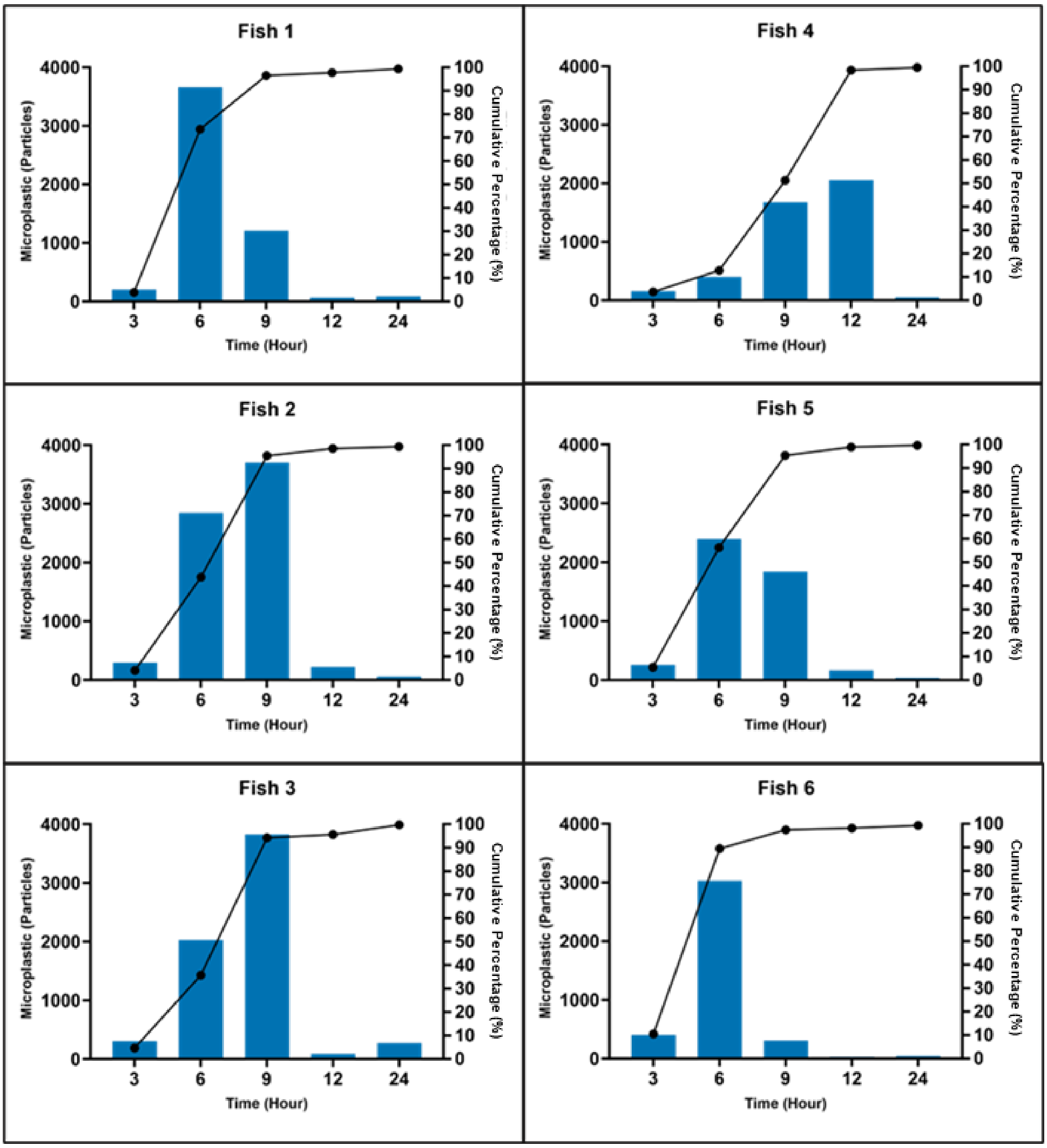


**Supplementary Figure S4. Microplastic elimination and timing in feeding *Oryzias javanicus* larvae in seawater.** Each bar shows numbers of eliminated MPs at each time point, whereas the line represents the cumulative percentage of MPs eliminated.

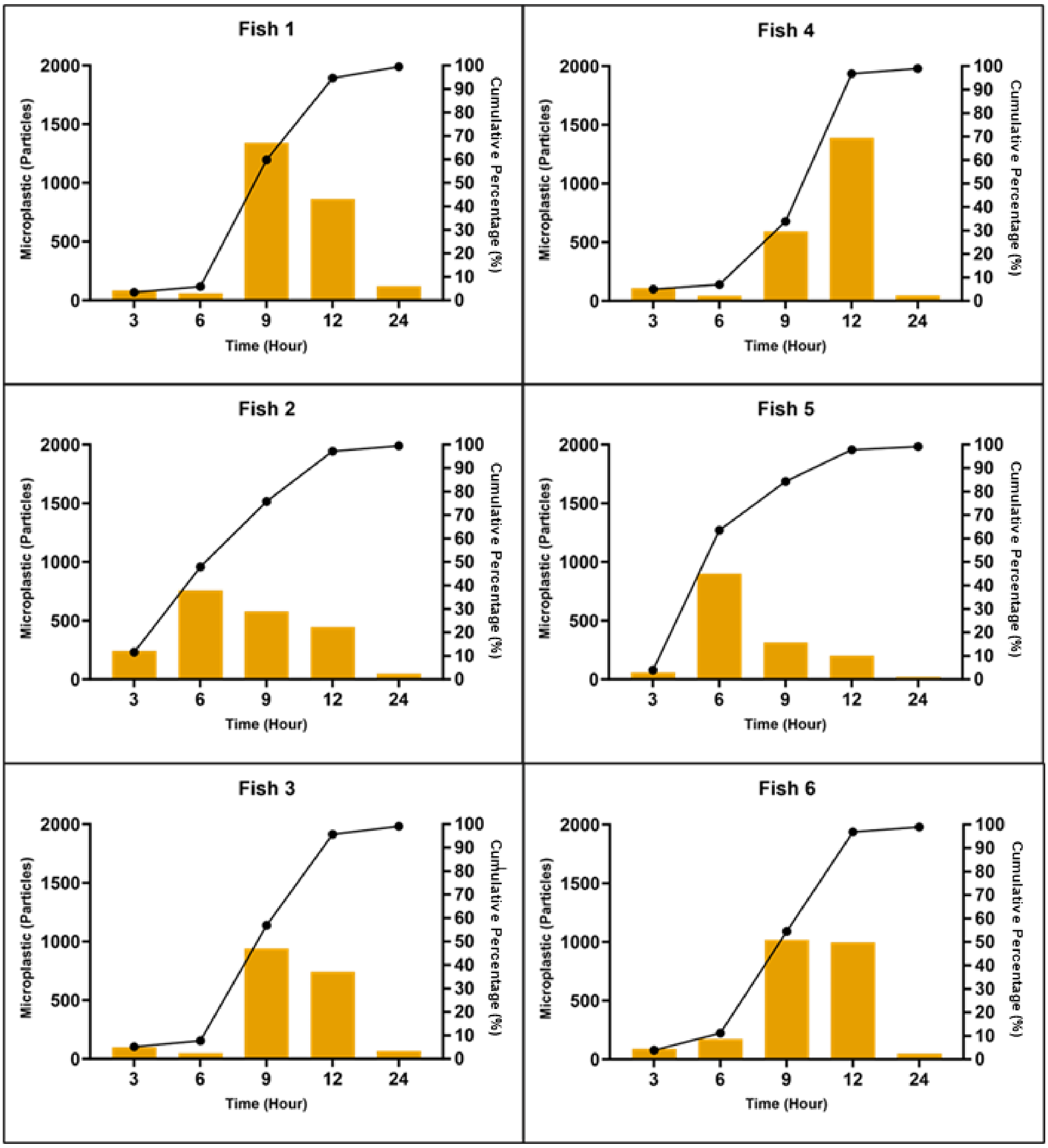


**Supplementary Figure S5. Microplastic elimination and timing in feeding *Oryzias javanicus* larvae in freshwater.** Each bar shows numbers of eliminated MPs at each time point, whereas the line represents the cumulative percentage of MPs eliminated.

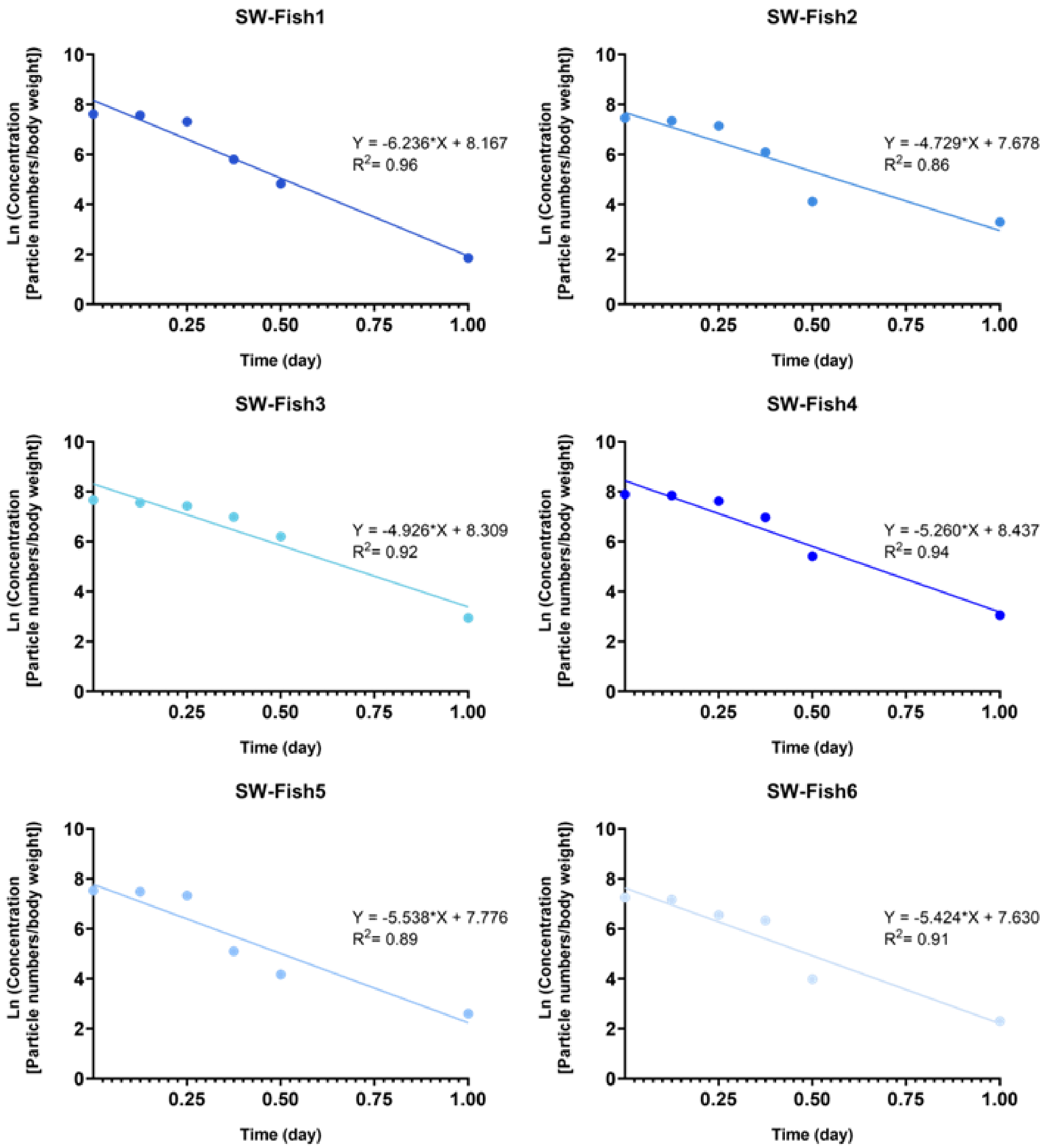


**Supplementary Figure S6. Linear regression model of microplastic elimination in each *Oryzias javanicus* larva in the seawater non-feeding group.** The linear elimination equation used for gut retention time calculations is written above the regression line. $R^2$ denotes the R squared value of each regression line.

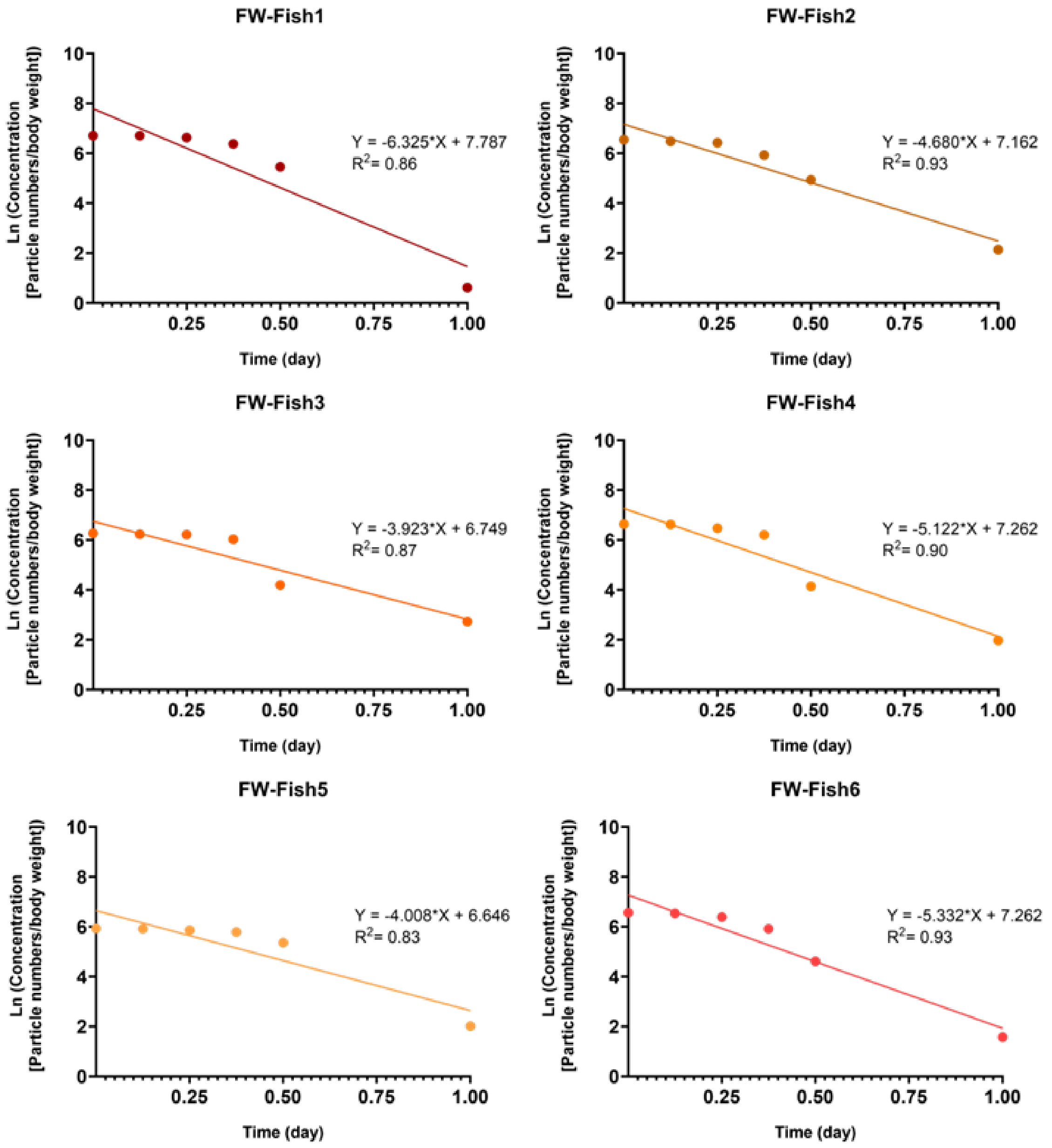


**Supplementary Figure S7. Linear regression model of microplastic elimination in individual *Oryzias javanicus* larva in the freshwater non-feeding group**. The linear elimination equation used for gut retention time calculations is written above the regression line. $R^2$ denotes the R squared value of each regression line.

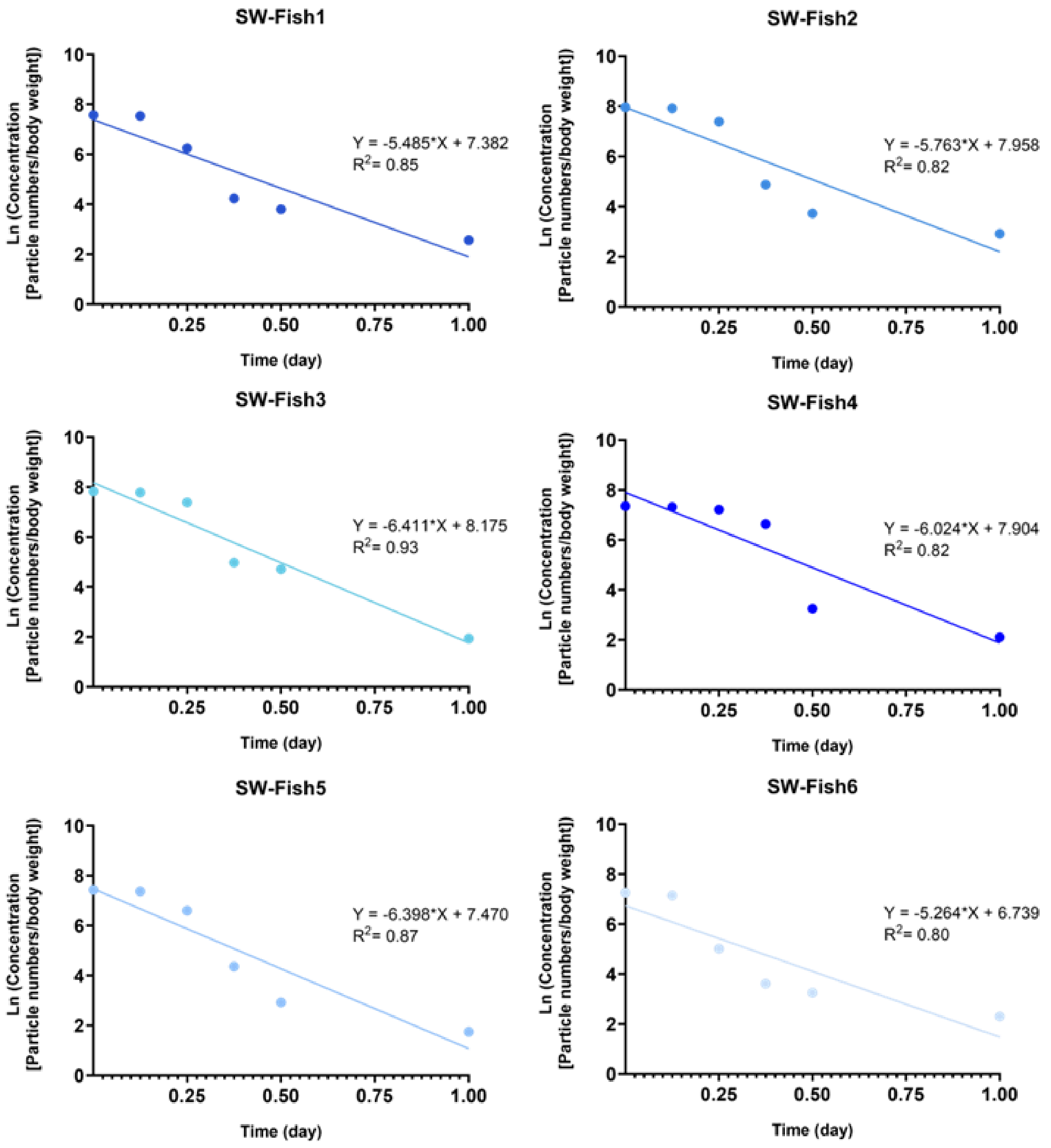


**Supplementary Figure S8. Linear regression model of microplastic elimination in individual *Oryzias javanicus* larva in the seawater feeding group.** The linear elimination equation used for gut retention time calculations is written above the regression line. $R^2$ denotes the R squared value of each regression line.

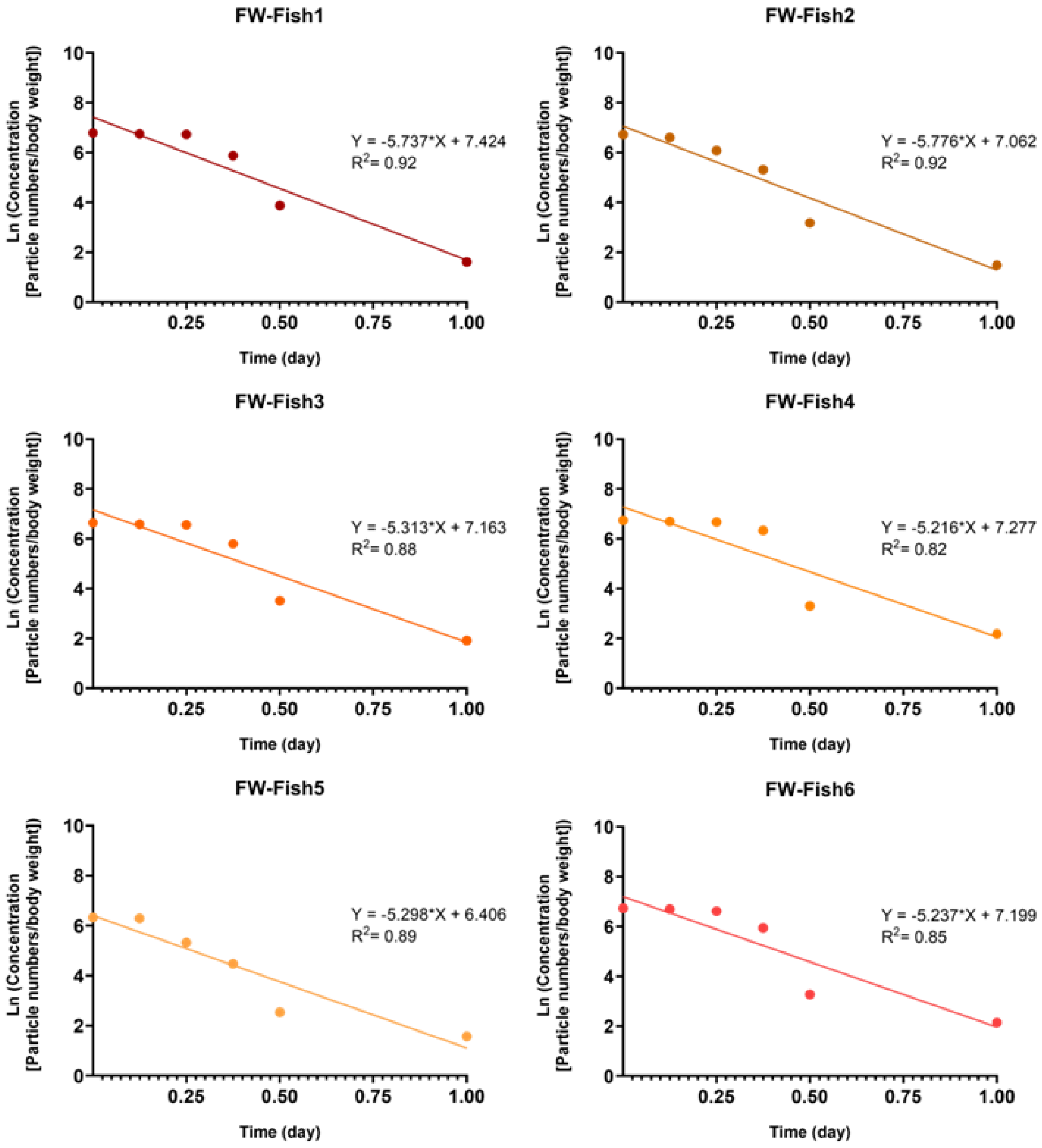


**Supplementary Figure S9. Linear regression model of microplastic elimination in individual *Oryzias javanicus* larva in the freshwater feeding group.** The linear elimination equation used for gut retention time calculations is written above the regression line. $R^2$ denotes the R squared value of each regression line.

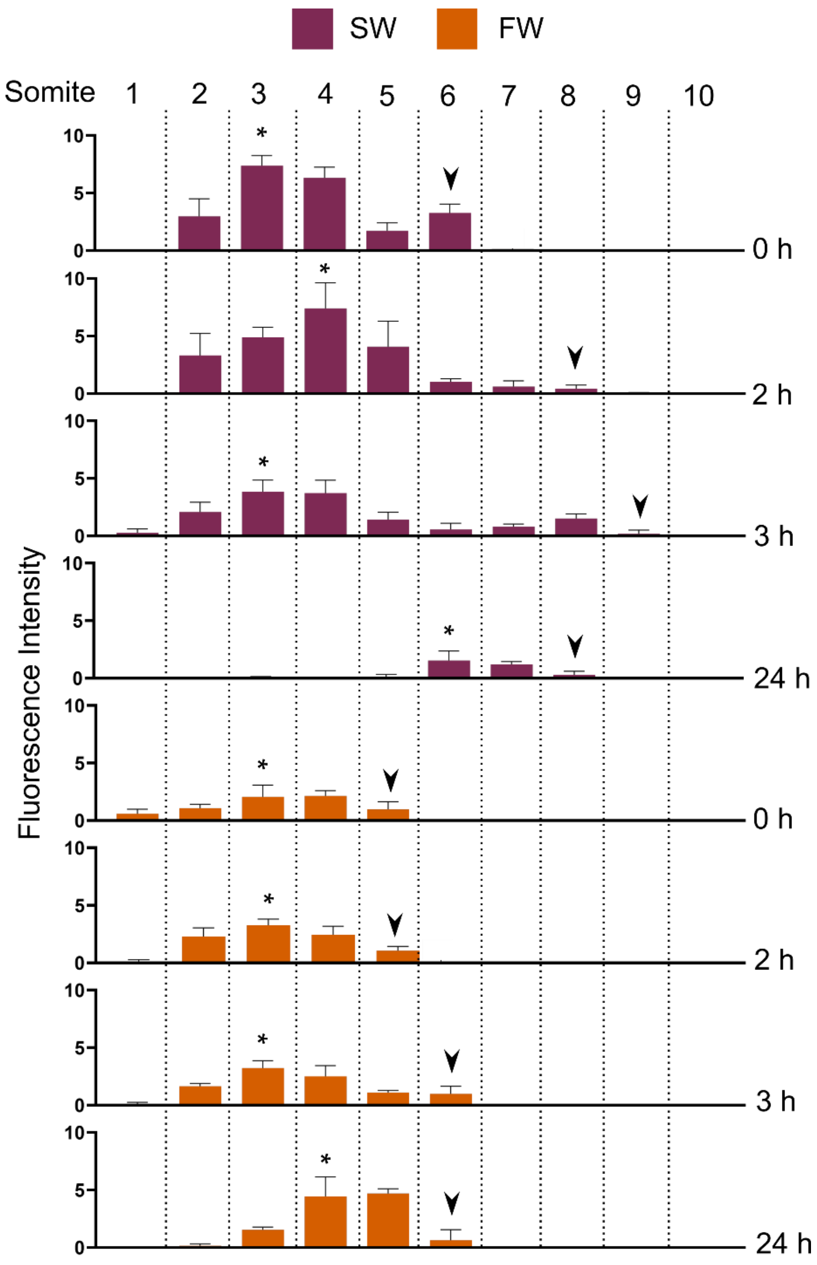


**Supplementary Figure S10. Comparison of FITC-dextran fluorescence intensity at each somite segment in SW and FW larvae.** Each bar shows the mean of FITC-dextran fluorescence intensity ± standard error of the mean (SEM) at each somite segment. Asterisks represents signal peaks, and arrowheads represent the most posterior part of the signal in the gastrointestinal tract.

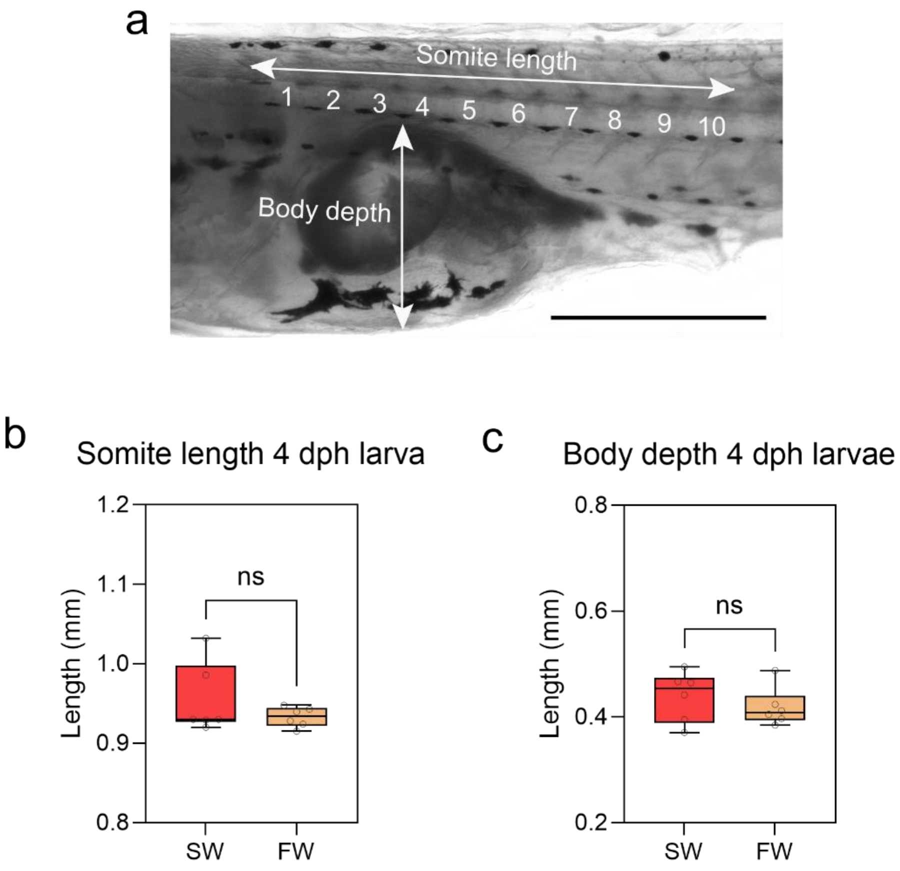


**Supplementary Figure S11. Measurement of somite length and body depth in 4 day-post hatching *Oryzias javanicus* larvae.** (a) Somite position in 4 dph *O. javanicus* larva, Comparison of (b) somite length and (c) Body depth of SW and FW larvae. Each bar shows the mean ± standard error of the mean (SEM). Differences between groups were analyzed for significance with Student's t-test (ns, non-significant $P > 0.05$).

**Supplementary Table S1. Water conditions during microplastic exposure and elimination in non-feeding groups.**

| Exposure Group | | Temperature (℃) | Salinity (ppt) | Density (kg/m³) | Viscosity (Pa•s) |
|---|---|---|---|---|---|
| **Seawater-reared *Oryzias javanicus* (without food)** | Exposure test | 26 | 31 | 1020.06 | 0.000929 |
| | Short-term elimination (0 h) | 26 | 31 | 1020.06 | 0.000929 |
| | Short-term elimination (3 h) | 26 | 31 | 1020.06 | 0.000929 |
| | Short-term elimination (6 h) | 25.9 | 31 | 1020.10 | 0.000931 |
| | Short-term elimination (9 h) | 25.8 | 31 | 1020.12 | 0.000934 |
| | Short-term elimination (12 h) | 25.9 | 31 | 1020.10 | 0.000931 |
| | Short-term elimination (24 h)<br>Long-term elimination (Day 0) | 25.9 | 31 | 1020.10 | 0.000931 |
| | Long-term elimination (Day 1) | 25.8 | 31 | 1020.12 | 0.000934 |
| | Long-term elimination (Day 2) | 25.9 | 31 | 1020.10 | 0.000931 |
| | Long-term elimination (Day 3) | 26 | 31 | 1020.06 | 0.000929 |
| | Long-term elimination (Day 4) | 25.9 | 31 | 1020.10 | 0.000931 |
| | Long-term elimination (Day 5) | 26 | 31 | 1020.06 | 0.000929 |
| **Freshwater-reared *Oryzias javanicus* (without food)** | Exposure test | 26 | 0 | 996.80 | 0.000870 |
| | Short-term elimination (0 h) | 25.9 | 0 | 996.81 | 0.000872 |
| | Short-term elimination (3 h) | 25.9 | 0 | 996.81 | 0.000872 |
| | Short-term elimination (6 h) | 25.8 | 0 | 996.88 | 0.000874 |
| | Short-term elimination (9 h) | 25.9 | 0 | 996.81 | 0.000872 |
| | Short-term elimination (12 h) | 26 | 0 | 996.80 | 0.000870 |
| | Short-term elimination (24 h)<br>Long-term elimination (Day 0) | 25.9 | 0 | 996.81 | 0.000872 |
| | Long-term elimination (Day 1) | 26 | 0 | 996.80 | 0.000870 |
| | Long-term elimination (Day 2) | 25.9 | 0 | 996.81 | 0.000872 |
| | Long-term elimination (Day 3) | 25.9 | 0 | 996.81 | 0.000872 |
| | Long-term elimination (Day 4) | 26 | 0 | 996.80 | 0.000870 |
| | Long-term elimination (Day 5) | 26 | 0 | 996.80 | 0.000870 |

*Temperatures and salinity were measured about 30 min after water replacement.

**Supplementary Table S2. Water conditions during microplastic exposure and elimination in feeding groups.**

| Exposure Group | | Temperature (℃) | Salinity (ppt) | Density (kg/m$^3$) | Viscosity (Pa•s) |
|---|---|---|---|---|---|
| **Seawater-reared *Oryzias javanicus* (with food)** | Exposure test | 25.9 | 31 | 1020.10 | 0.000931 |
| | Short-term elimination (0 h) | 26 | 31 | 1020.06 | 0.000929 |
| | Short-term elimination (3 h) | 25.9 | 31 | 1020.10 | 0.000931 |
| | Short-term elimination (6 h) | 25.9 | 31 | 1020.10 | 0.000931 |
| | Short-term elimination (9 h) | 26 | 31 | 1020.06 | 0.000929 |
| | Short-term elimination (12 h) | 25.9 | 31 | 1020.10 | 0.000931 |
| | Short-term elimination (24 h)<br>Long-term elimination (Day 0) | 26 | 31 | 1020.06 | 0.000929 |
| | Long-term elimination (Day 1) | 25.9 | 31 | 1020.10 | 0.000931 |
| | Long-term elimination (Day 2) | 25.8 | 31 | 1020.12 | 0.000934 |
| | Long-term elimination (Day 3) | 25.9 | 31 | 1020.10 | 0.000931 |
| | Long-term elimination (Day 4) | 26 | 31 | 1020.06 | 0.000929 |
| | Long-term elimination (Day 5) | 26 | 31 | 1020.06 | 0.000929 |
| **Freshwater-reared *Oryzias javanicus* (with food)** | Exposure test | 25.9 | 0 | 996.81 | 0.000872 |
| | Short-term elimination (0 h) | 26 | 0 | 996.80 | 0.000870 |
| | Short-term elimination (3 h) | 25.8 | 0 | 996.88 | 0.000874 |
| | Short-term elimination (6 h) | 25.8 | 0 | 996.88 | 0.000874 |
| | Short-term elimination (9 h) | 26 | 0 | 996.80 | 0.000870 |
| | Short-term elimination (12 h) | 26 | 0 | 996.80 | 0.000870 |
| | Short-term elimination (24 h)<br>Long-term elimination (Day 0) | 26 | 0 | 996.80 | 0.000870 |
| | Long-term elimination (Day 1) | 25.8 | 0 | 996.88 | 0.000874 |
| | Long-term elimination (Day 2) | 25.9 | 0 | 996.81 | 0.000872 |
| | Long-term elimination (Day 3) | 26 | 0 | 996.80 | 0.000870 |
| | Long-term elimination (Day 4) | 26 | 0 | 996.80 | 0.000870 |
| | Long-term elimination (Day 5) | 26 | 0 | 996.80 | 0.000870 |

*Temperatures and salinity were measured about 30 min after water replacement.

**Supplementary Table S3. Comparison of microplastic halftime ($T_{50}$) and elimination rate constant ($k_2$) in various fishes exposed to different polymers and sizes**

| Fish | | | MP Size (μm) | Feeding Status | $T_{50}$ (hour) | $k_2$ (day$^{-1}$) | References |
|---|---|---|---|---|---|---|---|
| Species | Stage | Habitat | | | | | |
| *Pagrus major* | Juvenile | SW | 250-300 | yes | 4.7 | 3.5 | Ohkubo et al., 2020 |
| *Fundulus heteroclitus* | Juvenile | SW | 250-301 | yes | 1.5 | 10.77 | Ohkubo et al., 2020 |
| *Fundulus heteroclitus* | Juvenile | SW | 710-850 | yes | 1.3 | 13 | Ohkubo et al., 2020 |
| *Carassius auratus* | Adult | FW | 50-500 | yes | 10 | - | Grigorakis et al., 2017 |
| *Oryzias latipes* | Juvenile | FW | 200 | yes | 14.8 | 1.13 | Liu et al., 2021 |
| *Oryzias latipes* | Juvenile | FW | 20 | yes | 16 | 1.04 | Liu et al, 2021 |
| *Oryzias latipes* | Juvenile | FW | 20 | yes | 17.7 | 0.94 | Liu et al., 2021 |
| *Oryzias latipes* | Juvenile | FW | 2 | yes | 21.8 | 0.76 | Liu et al., 2021 |
| *Oncorhynchus mykiss* | Adult | FW | 42.7 | yes | 12.1 | - | Roch et al., 2021 |
| *Oncorhynchus mykiss* | Adult | FW | 1,086 | yes | 4 | - | Roch et al., 2021 |
| *Cyprinus carpio* | Adult | FW | 42.7 | yes | 7.3 | - | Roch et al., 2021 |
| *Cyprinus carpio* | Adult | FW | 1,086 | yes | 4.6 | - | Roch et al., 2021 |
| *Oryzias javanicus* | Larva | SW | 1 | yes | 3.5 | 5.527 | This study |
| *Oryzias javanicus* | Larva | SW | 1 | no | 5.6 | 5.198 | This study |
| *Oryzias javanicus* | Larva | FW | 1 | yes | 5.2 | 5.406 | This study |
| *Oryzias javanicus* | Larva | FW | 1 | no | 7.0 | 4.529 | This study |

-; not mentioned in the article
